# Microbial resource management for *ex situ* biomethanation of hydrogen at alkaline pH

**DOI:** 10.1101/2020.03.18.995811

**Authors:** Washington Logroño, Denny Popp, Sabine Kleinsteuber, Heike Sträuber, Hauke Harms, Marcell Nikolausz

**Author notes:** Correspondence; (M.N.).

## Abstract

Biomethanation is a promising solution to convert H_2_ produced from surplus electricity and CO_2_ to CH_4_ by using hydrogenotrophic methanogens. In *ex situ* biomethanation with mixed cultures, homoacetogens and methanogens compete for H_2_/CO_2_. We enriched a hydrogenotrophic microbiota on CO_2_ and H_2_ as sole carbon and energy sources, respectively, to investigate these competing reactions. Microbial community structure and dynamics of bacteria and methanogenic archaea were evaluated through 16S rRNA and *mcrA* gene amplicon sequencing, respectively. Hydrogenotrophic methanogens and homoacetogens were enriched as acetate was concomitantly produced along with CH_4_. By controlling the media composition, especially changing the reducing agent, the formation of acetate was lowered and grid quality CH_4_ (≥ 97%) was obtained. Formate was identified as an intermediate that was produced and consumed during the bioprocess. Stirring intensities ≥1000 rpm were detrimental, probably due to shear force stress. The predominating methanogens belonged to the genera *Methanobacterium* and *Methanoculleus*. The bacterial community was dominated by *Lutispora*. The methanogenic community was stable, whereas the bacterial community was more dynamic. Our results suggest that hydrogenotrophic communities can be steered towards selective production of CH_4_ from H_2_/CO_2_ by adapting the media composition, the reducing agent and the stirring intensity.

## 1. Introduction

Renewable energy from wind power and photovoltaics increasingly leads to temporary excess of electricity that cannot be handled by the grid and traditional storage infrastructure. Hence, technical solutions to store this energy, e.g. in form of chemical energy carriers, are required. The power-to-gas (P2G) technology converts surplus power into a storable gas [1]. H_2_ can be generated through water electrolysis and subsequently injected and stored in the natural gas grid, though with certain limitations [2]. Methane can also be produced from excess electricity in a two-stage process: H_2_ is first produced through water electrolysis and then used in a methanation stage to reduce CO_2_ to methane [2]. Although the H_2_ production technology is quite advanced, it has some drawbacks concerning long-term storage, safety and low energy density of H_2_ as well as the requirement of technical modifications of the natural gas grid. Methane, on the other hand, is very attractive because the storage and distribution infrastructure is already in place in many countries. Methane can be readily injected into the gas grid and has a volumetric energy content of 36 MJ m^−3^, which is more than three times higher than that of H_2_ (10.88 MJ m^−3^) [3].

Biogas is the product of anaerobic digestion (AD), which is a well-established commercial process and a key technology in the current and future renewable energy sector [4]. Biogas consists mainly of CH_4_ (40-75%) and CO_2_ (25-60%) and needs to be upgraded to biomethane by removing CO_2_ if injection into the gas grid is intended. Methods for biogas upgrading have been reviewed elsewhere [5–7]. Biological biogas upgrading (biomethanation) uses external H_2_ to convert the CO_2_ share of the biogas into additional CH_4_ via the CO_2_-reductive pathway of hydrogenotrophic methanogens. Biomethanation of H_2_ is an emerging technology that appears to be advantageous over the catalyst-based chemical methanation (Sabatier reaction) in terms of milder reaction conditions [6]. This bioprocess can be performed by pure methanogenic strains [8] or mixed cultures [6]. The latter may have certain economic and process advantages over pure cultures [9].

According to Kougias et al. [10] and Rittmann [11], biomethanation of H_2_ can be done in three ways: *in situ*, *ex situ* and by a hybrid process. In the *in situ* process, H_2_ is injected into the main anaerobic digester or post digester of a biogas plant to reduce CO_2_ and thereby increase the CH_4_ content of the biogas. In the *ex situ* process, biogas or CO_2_ reacts with H_2_ in a bioreactor that is separate from the AD process. The hybrid process couples partial biogas upgrading in the main AD reactor (*in situ*) with a final upgrading step in a separate reactor (*ex situ*). Defining the system to be investigated according to the above mentioned categories is important for comparisons in terms of efficiency and microbiota. An *ex situ* reactor could provide a defined ecological niche to enrich specialized hydrogenotrophic microbiota with autotrophic metabolism (methanogenesis and homoacetogenesis). It can be hypothesized that the inoculum, the operation temperature and the continuous supply of H_2_ play important roles in shaping the microbial community towards the predominance of either hydrogenotrophic methanogens or homoacetogenic bacteria.

If a complex inoculum is used to perform *ex situ* biomethanation of H_2_, acetate could be synthesized from H_2_ and CO_2_ via the Wood-Ljungdahl pathway concomitantly with CH_4_ formation. Acetate synthesis during *ex situ* biomethanation represents a problem as it is an undesirable carbon and electron sink when CH_4_ is the target molecule. It is therefore necessary to manage the microbiota towards selective CH_4_ production. In environmental biotechnology, the term microbial resource management implies finding strategies to obtain and maintain a highly performing community [12]. The understanding of metabolic processes in complex communities imposes a great challenge that could be overcome by establishing enrichment cultures to investigate the essential metabolic functions. Enrichment cultures are used to construct mixed culture metabolic models and explore the ecophysiological functions of microbes that have not yet been isolated in pure culture [13]. Enriched mixed cultures could be simple enough to investigate individual community members [14] and represent opportunities to grow uncultivable microbes [15]. In fact, some microbes can only grow in the presence of other microorganisms [16] and this could be true for the microorganisms of the AD microbiota that still remains largely unexplored. In the *ex situ* H_2_ biomethanation context, most studies resulted in the enrichment of methanogens and homoacetogens. Previous *ex situ* biomethanation studies have found *Methanobacteriales* [10,17–23], *Methanomicrobiales* [10,24] and *Methanococcales* [24] as the dominant orders. *Methanothermobacter thermautotrophicus* was the dominant methanogen in three different reactor configurations [10]. In the bacterial domain, *Firmicutes* [10,21,25,26], *Bacteroidetes* [10], *Synergistetes* [21] and *Proteobacteria* [25] were the dominant phyla.

In previous studies, sludge from biogas reactors or wastewater anaerobic granules were used as inoculum sources [10,17,19,24,27–31]. From the microbiological point of view, certain process parameters such as temperature and pH, substrate characteristics and inoculum source define the community structure and dominant members of the microbiota. An *ex situ* study comparing different reactor configurations reached methane concentrations of more than 98%; however, acetic acid accumulated to concentrations of ~4 g L^−1^ and the pH values were ≥ 8 [10]. Similar pH values ranging from 7 to ≥ 8 were also reported in a continuous stirred tank reactor (CSTR) for *ex situ* biomethanation [32]. Previous *ex situ* biomethanation studies have operated at slightly alkaline pH, but microbial enrichment studies at such conditions are still missing.

In the present study, we explored *ex situ* biomethanation of H_2_ at alkaline pH through an enrichment process. The aim of the enrichment strategy was to better understand the microbial community dynamics depending on the operational conditions as well as the competition between methanogenesis and homoacetogenesis during *ex situ* biomethanation. According to previous studies, hydrogenotrophic methanogenesis is the major methanogenic pathway under high nitrogen load and high ammonia concentration [33–35]. Thus, we used the digestate of a laboratory-scale biogas reactor (CSTR) treating a nitrogen-rich substrate (dried distillers grains with solubles, DDGS) as inoculum source for long-term enrichment of a hydrogenotrophic microbiota that performs *ex situ* biomethanation. The experiments focused on the characterization of the microbial community structure and dynamics during the enrichment by amplicon sequencing of 16S rRNA and *mcrA* genes. Additionally, the effects of media components such as yeast extract and reducing agent as well as of the stirring intensity on the H_2_ and CO_2_ metabolism were investigated.

## 2. Materials and Methods

### 2.1 Inoculum

Anaerobic sludge from a mesophilic (38°C) laboratory-scale CSTR treating DDGS was sieved using a 400-μm mesh size sieve under nitrogen flow. The liquid inoculum was degassed at 38°C for 7 days before use. The basic characteristics are the mean values of triplicate measurements as follows: total solids (TS), 3.4%; volatile solids (VS), 70.1%_TS_; pH, 7.5.

### 2.2 Growth medium

Modified mineral medium DSMZ1036 containing yeast extract (0.2 g L^−1^) as described previously [36] was used for the enrichment and is designated as medium A in the following. For further experiments, the medium was used in two variants: medium B did not contain yeast extract but was supplemented with a vitamin solution as described by [37] and cysteine-HCl as reducing agent in the same concentration as in medium A. Medium C contained vitamins like medium B but sodium sulfide as reducing agent as described by [37]. After preparing the media as described in **Text S1**, the pH for all media variations was adjusted to 9 with a sterile anoxic stock solution of 2 M KOH.

### 2.3 Enrichment setup

Strict anaerobic techniques were thoroughly applied in this study. Sterile anoxic bottles were prepared as described in **Text S2** prior to medium dispensing and inoculation. The gas volume/liquid volume ratio was maintained at 3 for all experiments regardless of the size of the bottle, unless stated otherwise. The experiments were conducted with four biological replicates in the first stage (gas feeding of the anaerobic sludge) and triplicates in the second stage (enrichment in mineral medium). The detailed chronology of the culture transfers is provided in **Table S4.**

The setup in the first stage was assembled in an anaerobic chamber. Serum bottles of 219.5 mL volume were filled with 50 mL degassed inoculum, sealed with butyl rubber stoppers and crimped with aluminum caps. The gas phase of the serum bottles was replaced by H_2_ (80%) and CO_2_ (20%). All bottles receiving H_2_ and CO_2_ were operated in fed-batch mode and daily pressurized to ~2.2 bar for approximately five months. Bottles containing the inoculum and a nitrogen atmosphere (not pressurized) were used as controls to account for the residual biogas production. Detailed information about headspace flushing and pressurization is given in **Text S2**.

In the second stage, medium A was used to enrich a particle-free culture by six subsequent culture transfers in fresh medium bottles by inoculating the content of the previous culture transfer (10%, v/v). One randomly selected replicate from the first stage served as inoculum to start the bottles of the second stage. Anoxic medium A (45 mL) was dispensed to sterile, anoxic serum bottles and left overnight in an incubator at 37°C to reduce any oxygen traces that entered the bottles during medium dispensing. Next, the bottles were inoculated with 5 mL culture from the first stage. Biological controls for determining residual biogas production (containing inoculum but with N_2_ gas phase) as well as sterile controls (not inoculated, but with either H_2_/CO_2_ or N_2_ gas phase) were also set up. The bottles were fed with gaseous substrate as described above and incubated at 37.4°C in an orbital shaking incubator (IKA KS 4000 ic control, IKA®-Werke GmbH & Co. KG) at 200 rpm.

### 2.4 Cultivation experiments

In a series of four independent experiments, the effects of medium composition and stirring intensity on biomethanation, biomass growth, and production of volatile fatty acids (VFA) were investigated. Experiments were conducted in 1-L pressure-resistant Duran bottles (Schott AG, Germany). Bottles were inoculated with 10% (v/v) pre-culture (21 days old second stage enrichment culture, 11^th^ transfer (T11)) and incubated at 37.4°C under constant orbital shaking at 200 rpm. The experiments were conducted in duplicates and stepwise to investigate the effect of the medium composition (media A, B and C as described in section 2.2). The gas consumption and production as well as the development of biomass and VFA production were frequently monitored.

After the optimal medium had been determined, we tested the effect of the stirring intensity on the CH_4_ production and H_2_/CO_2_ consumption. Instead of shaking, the bottles were stirred with magnetic stirrers (top plate diameter of 145 mm, speed range from 100 to 1,400 rpm; Heidolph, Germany) and new magnetic bars (50 mm × 8 mm, LABSOLUTE, Th. Geyer, Germany) and incubated at 38°C. To reduce detrimental effects of shear forces on the cells, the liquid volume was increased to 500 mL, corresponding to a gas volume/liquid volume ratio of 1. The experiments were conducted three times with duplicates (n=6).

Flushing and pressurizing as well as pressure determination and sampling of the gas and liquid phases were done as described above (see also **Text S2**) for all four experiments.

### 2.5 Microbial community analysis

Samples for community analysis were taken from the inoculum (S) as well as after one month (1M) and 5 months (5M) fed-batch feeding during the first stage of the enrichment. Samples from the second stage were taken at the end of the first (T1), third (T3), and sixth (T6) culture transfer. Liquid samples of 1.5 mL were withdrawn from each bottle with a nitrogen-flushed syringe (**Text S2**) and centrifuged at 4°C and 20,817 × *g* for 10 min. Pellets were stored at −20°C until DNA extraction. DNA was extracted with the NucleoSpin®Soil Kit (MACHEREY-NAGEL GmbH & Co. KG, Germany) using buffer SL2 and enhancer solution. The quality and quantity of extracted DNA were verified via gel electrophoresis (0.8% agarose) and photometrically using a NanoDrop ND 1000 spectral photometer (Thermo Fisher Scientific, USA). Extracted DNA was stored at −20°C until use. The microbial community composition was analyzed by amplicon sequencing of *mcrA* genes for methanogens and 16S rRNA genes for bacteria.

In order to analyze the bacterial communities, the V3-V4 region of the 16S rRNA genes was amplified using the universal primers 341f (5’-CCT ACG GGN GGC WGC AG-3’) and 785r (5’-GAC TAC HVG GGT ATC TAA KCC-3’) described by Klindworth et al. [38]. For the analysis of the methanogenic communities, the mlas (5’-GGT GGT GTM GGD TTC ACM CAR TA-3’) and mcrA-rev (5‘-CGT TCA TBG CGT AGT TVG GRT AGT-3’) primers were used as described previously [39]. All primers contained Illumina MiSeq-specific overhangs. Amplicon libraries were prepared and sequenced on the Illumina MiSeq platform using the MiSeq Reagent Kit v3 with 2×300 cycles. Demultiplexed raw sequence data were deposited at the EMBL ENA under the study accession number PRJEB36972 (http://www.ebi.ac.uk/ena/data/view/PRJEB36972).

Primer sequences were clipped from demultiplexed and adapter-free reads using Cutadapt v1.18 [40]. Further sequence analysis was performed using QIIME2 v2019.1 [41]. Sequences were trimmed, denoised and merged using the dada2 plugin [42]. For 16S rRNA gene analysis, forward and reverse reads were truncated at 270 bp and 240 bp, respectively. For *mcrA* gene analysis, reads were truncated at 270 bp and 230 bp, respectively. Maximum expected errors were set to 2, which is the default value. Chimeras were removed in default consensus mode of the dada2 plugin. Resulting feature sequences of 16S rRNA gene analysis were classified against the MiDAS database v2.1 [43] trimmed to the region covered by the 341f and 785r primers. For *mcrA* gene analysis, a taxonomy database was created by using *mcrA* sequences from the RDP FunGene database [44]. For this purpose, *mcrA* sequences were downloaded (2,878 sequences on 21^st^ January 2019), sequences from uncultured organisms and metagenomic sequences were removed, and taxonomic information was formatted resulting in 385 sequences used for the classification. As the primer combination 341f/785r also amplifies archaeal 16S rRNA genes, archaeal reads were removed from further analysis of the 16S rRNA genes and bacterial read counts were normalized to 100%.

### 2.6 Analytical methods

To determine the TS content of the inoculum, samples were dried at 105°C for 24 h and the mass was recorded. The TS value was calculated from the mass difference of the fresh and dried sample. Subsequently, the samples were incinerated at 550°C in a muffle furnace for 2 h and the mass was recorded. The VS value was calculated based on mass difference of dried and incinerated samples. The mean values of triplicate measurements are presented.

To determine the headspace gas composition, 1 mL gas sample was withdrawn with a syringe and injected into an argon pre-flushed glass vial of 20 mL (**Text S2**). The gas samples were measured via gas chromatography equipped with an autosampler in a Perkin Elmer GC. The GC was equipped with HayeSep N / Mole Sieve 13X columns and a thermal conductivity detector. The oven and detector temperatures were 60°C and 200°C, respectively. The carrier gas was argon. Every gas sample was analyzed immediately or within 24 h after sampling.

The relative pressure in the bottles was measured with a digital manometer (**Text S2**). The gas amount in the bottles was calculated according to Equation 1 (Eq. 1). Standard conditions were considered for calculations (P = 1.01325 bar, T = 298.15 K). The consumption and production rates of gases (H_2_, CO_2_, and CH_4_) were determined from the linear slope of at least three continuous measurements and are given in mmol gas per liter liquid volume per hour (mmol L^−1^ h^−1^).

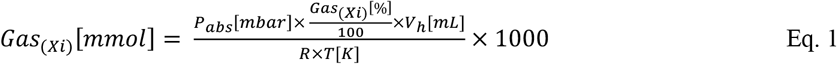

Where, Xi refers to the gas in question, P_abs_ is the absolute pressure inside the bottle, V_h_ is the headspace volume of the cultivation bottle (169.5 mL), R is the universal gas constant (8.314 × 10^4^ mbar cm^3^ mol^−1^ K^−1^), and T is the standard temperature.

For measuring the concentration of VFA (formic, acetic, propionic and butyric acid), the supernatants from liquid samples were filtered through a membrane filter with 0.2 μm pore size (13 mm; LABSOLUTE, Th. Geyer GmbH, Germany) and stored at −20°C or analyzed immediately. When needed, appropriate dilutions were prepared with deionized water and the samples were analyzed by using high performance liquid chromatography (HPLC; Shimadzu Scientific Instruments, US) equipped with an RID detector L-2490 and an ICSep column COREGEL87H3 (Transgenomic Inc., USA). The sample volume for HPLC measurement was 200 μL and the injection volume was 20 μL. The HPLC measurements were done with 5 mM H_2_SO_4_ as eluent at a flow rate of 0.7 mL min^−1.^

The pH value of the broth was measured in 200 μL liquid samples using a mini-pH meter (ISFET pH meter S2K922, ISFETCOM Co., Ltd., Japan) and the value was recorded after 90 s. For particular experiments with medium B and C, the pH was determined as aforementioned after a centrifugation step at 20,817 × *g*, 4°C, 10 min.

### 2.7 Statistical analysis

The methane concentrations of the two stages were compared when the gas conversion and production was stable throughout several feeding cycles (variation was less than 10% in at least ten consecutive batch-cycle feedings) by means of analysis of variance (ANOVA). A Tukey post-hoc test was used for multiple comparisons with a confidence level of 0.05. RStudio [45], Graphpad (Graphpad Software, Inc., San Diego, CA) or Microsoft Excel were used to compute the data. Microbial community composition data were analyzed by principal coordinate analysis (PCoA) based on Bray-Curtis distances of relative abundances in addition to the absence and presence using the phyloseq package [41] version 1.30.0 in R [46] version 3.6.1. PCoA was plotted using the ggplot2 package [47] version 3.2.1.

## 3. Results and Discussion

### 3.1 Enrichment of the hydrogenotrophic community and biomethanation performance

During the first stage of the enrichment, methane was formed within 24 h upon H_2_/CO_2_ feeding, and the process was stable for ~5 months. The rapid gas substrate conversion indicated high hydrogenotrophic methanogenesis activity in the inoculum. This observation is in agreement with a previous study [48]. However, the highest methane concentration between feeding cycles was only ~90% (**Figure S1**). In the second stage of the enrichment, the CH_4_ concentrations in six successive transfers (T1-T6) were as high as in the bioreactors working with sludge in the first stage (**Table 1**). The CH_4_ concentration was 6% lower than the one described in a previous study [17] but similar to that observed in another previous study [20].

**Table 1.** Summary of process parameters during the enrichment in the first stage with sludge and in the second stage with six culture transfers (T1-T6).

| Sample | Sludge <sup>b</sup> | T1 <sup>c</sup> | T2 <sup>c</sup> | T3 <sup>c</sup> | T4 <sup>c</sup> | T5 <sup>c</sup> | T6 <sup>c</sup> |
| --- | --- | --- | --- | --- | --- | --- | --- |
| Days of incubation | 167 | 56 | 27 | 22 | 24 | 28 | 40 |
| CH<sub>4</sub> (%)<sup>a</sup> | 85.17 ± 4.5 | 86.99 ± 5.4 | 88.04 ± 3.2 | 87.48 ± 4.7 | 88.00 ± 2.3 | 87.21 ± 4.1 | 87.47 ± 2.5 |
| Biomass concentration<br>start (mg L<sup>-1</sup>)<sup>d</sup> | - | - | - | 92.2 ± 11.73 | 91.4 ± 2.56 | 90.3 ± 17.79 | 88.6 ± 10.48 |
| end (mg L<sup>-1</sup>)<sup>d</sup> | - | - | - | 609.1 ± 33.57 | 548.2 ± 30.22 | 570.7 ± 31.46 | 588.2 ± 32.42 |
| pH (start) | 8.0 ± 0.1 | 9.0 ± 0.1 | 9.0 ± 0.1 | 9.0 ± 0.1 | 9.0 ± 0.1 | 9.0 ± 0.1 | 9.0 ± 0.1 |
| pH (end) | 8.5 ± 0.2 | 8.4 ± 0.1 | 8.3 ± 0.2 | 7.9 ± 0.1 | 8.1 ± 0.1 | 8.0 ± 0.1 | 8.1 ± 0.1 |
| Acetate (mg L<sup>-1</sup>) | 123.5 ± 7.8 | 2089 ± 194 | 970.8 ± 54.6 | 1922 ± 398 | 2053 ± 156 | 886.9 ± 170.9 | 122.7 ± 20.1 |
| Propionate (mg L<sup>-1</sup>) | 0.0 ± 0.0 | 26.03 ± 2.3 | 492.1 ± 16.4 | 185.9 ± 18.3 | 188.3 ± 37.9 | 135.9 ± 3.9 | 92.15 ± 3.7 |
| Butyrate (mg L<sup>-1</sup>) | 0.0 ± 0.0 | 56.33 ± 19.2 | 0.0 ± 0.0 | 147.2 ± 13.2 | 84.04 ± 10.26 | 28.1 ± 0.84 | 25.89 ± 1.4 |

<sup>a</sup> Concentration measured when the gas production was stable (mean and standard deviation, n=17).
<sup>b</sup> Four biological replicates. Mean and standard deviation is shown.
<sup>c</sup> Three biological replicates. Mean and standard deviation is shown
<sup>d</sup> Experimental conversion factor for biomass quantification: 1 OD<sub>600</sub> = 0.423 g L<sup>-1</sup> dry weight. Details are given in **Figure S3**.

Particle-free cultures were obtained after the third transfer and therefore the cell biomass from T3-T6 could be followed by spectrophotometry via optical density (OD_600_) (**Figure S2**). On average, each transfer from T3 to T6 started with a biomass of 91 ± 22 mg L^−1^ whereas the final biomass was on average 579 ± 26 mg L^−1.^ In terms of biomass and gas composition no significant difference (p ≤ 0.05) among transfers was found when comparing the end points of each transfer (**Table 1**). At the end of each culture transfer, the pH was ~8, a value similar to that reported in a previous study [10]. Acetate was found in considerable concentrations from T1 to T5 but not in the seed sludge (after ~5 months) and in T6 (see **Table 1**), indicating that homoacetogenesis was a concomitant reaction along with methanogenesis in our enriched hydrogenotrophic community. The variation in acetate concentrations among transfers could be associated with the cultivation time, especially for the particle-free enrichment cultures (T5 and T6), where the acetate concentration was generally low and the cultivation time was long.

The decrease of the acetate concentration at the end of the culture transfer (T6) suggests decreased homoacetogenic activity or increased acetate utilization via syntrophic acetate oxidation (SAO) since acetotrophic methanogens were absent in our enrichment cultures (see section 3.2). Another explanation for the decrease could be acetate assimilation to build up microbial biomass since hydrogenotrophic microbes can use an organic carbon source such as acetate when available. It was previously reported that acetate is central to the carbon metabolism of autotrophic and heterotrophic microbes [49]. Despite the variations of acetate concentration, the observed values throughout the culture transfers were similar to those reported in other studies [10,19,24,26,30]. Since acetate was the main side product, the kinetics of acetate consumption was assessed in more detail during one batch-cycle feeding when the culture (T1) presented stable gas consumption in several consecutive batch cycles (**Figure S3**). Acetate was consumed during the first 7 h (4% consumption likely to build biomass) and then its concentration increased again (1% increase), although the final concentration was 3% less than the initial concentration. Propionate and butyrate were also detected and the highest concentrations were found in T1 and T3. However, their concentrations decreased over the consecutive transfers (**Table 1**). Most likely this effect was also related to the cultivation time of the culture transfers.

### 3.2 Microbial community structure and dynamics

Feeding the complex community with a rather simple substrate has probably reduced the microbial diversity as shown previously [10] but it could still maintain a number of cooperative as well as competing functional groups. The effect of H_2_/CO_2_ as selection factor shaping the bacterial and methanogenic communities resulted in a dynamic process throughout the enrichment as visualized by PCoA (see time trajectory in **Figure 1**). The first two axes explained 64% and 98% of the variance for the bacterial (**Figure 1A**) and methanogenic (**Figure 1B**) communities, respectively. Hence, a two-dimensional plot is sufficient to represent the relation between the samples. Microbial communities grouped according to transfers, which means that the communities of the same transfer were quite similar but different from those of the other transfers.

**Figure 1.**
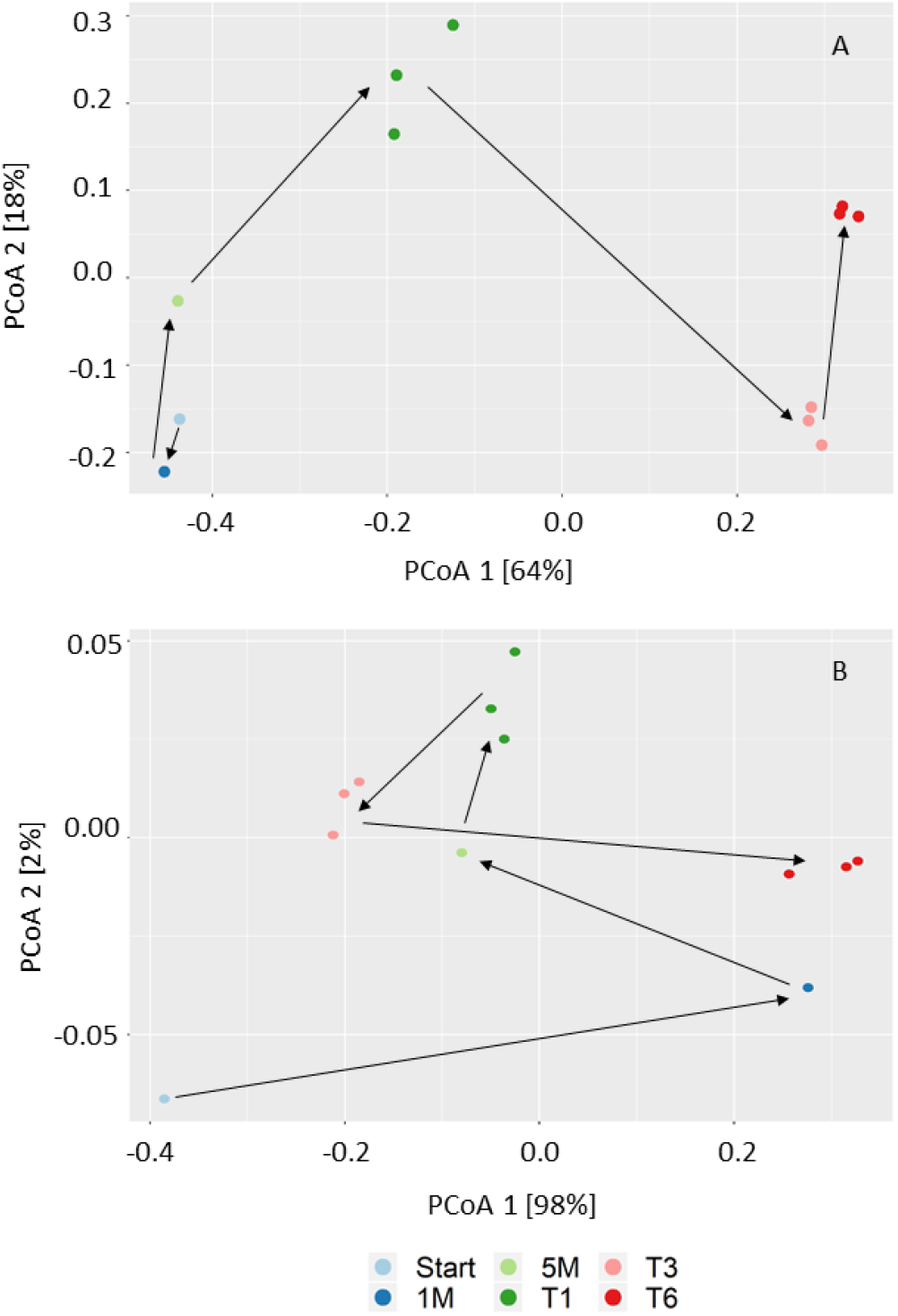
Principal coordinate analysis (PCoA) based on Bray-Curtis distances showing the microbial community shift during the enrichment. A) Bacterial community (16S rRNA gene amplicon sequences) and B) methanogenic community (*mcrA* gene amplicon sequences). The labels of the figure are as follows: inoculum (Start), one month (1M), 5 months (5M) after fed-batch feeding during the first stage; first transfer (T1), third transfer (T3), and sixth transfer (T6) after fed-batch feeding in the second stage.

As expected based on the high ammonia level (5.84 g L^−1^ NH_4_-N) of the source digester, the methanogenic community in the inoculum was dominated by hydrogenotrophic methanogens affiliated to the genus *Methanobacterium* (**Figure 2**), which is in agreement with previous studies on high ammonia level reactors [33–35]. Methanogens affiliated to the genus *Methanosarcina* were also present in the inoculum but disappeared after one month of H_2_/CO_2_ feeding despite their versatility in substrate utilization. It was proposed that H_2_ feeding exerts a selection pressure to enrich hydrogenotrophic methanogens [21,25], which could explain the disappearance of *Methanosarcina* and the complete dominance of strictly hydrogenotrophic methanogens in the enrichment. Furthermore, the H_2_ uptake rate was reported to be one order of magnitude higher for the strict hydrogenotrophic methanogen *Methanococcus maripaludis* [50] than for the versatile methanogen *Methanosarcina barkeri* [51], thus considering this aspect may also explain why *Methanosarcina* disappeared although it can grow on H_2_/CO_2_.

**Figure 2.**
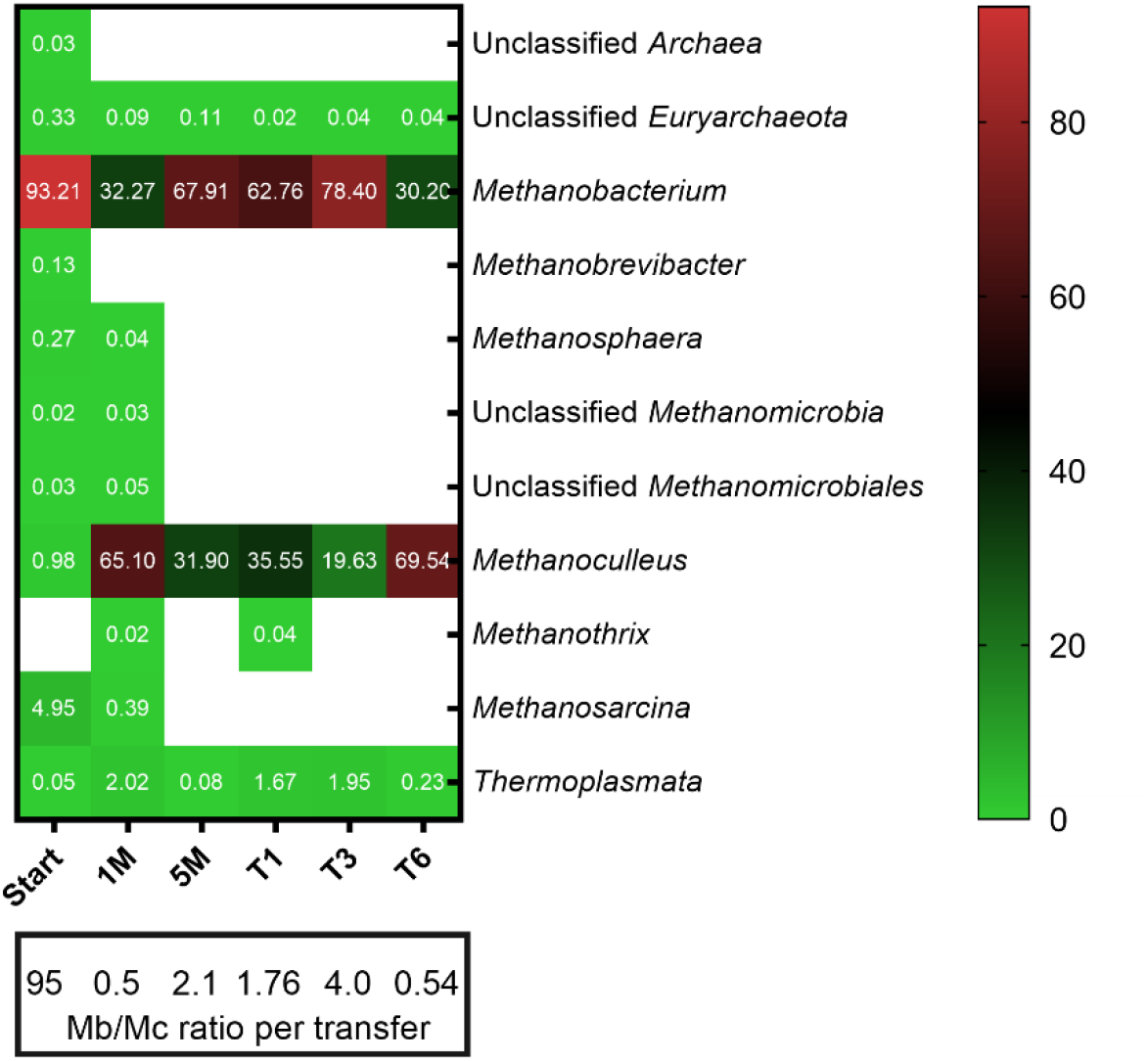
Methanogenic community structure in different stages of the enrichment. Taxa with a relative abundance less than 0.01% were filtered out from all samples. Numbers represent the relative abundance in percent and blank space indicates the absence of the respective taxa. The ratio of the most dominant methanogens *Methanobacterium* (Mb) and *Methanoculleus* (Mc) among transfers is shown. Mean values of three biological replicates are presented for T1, T3 and T6 whereas single values are shown for Start, 1M and 5M.

Species of the genus *Methanoculleus* dominated (65% of the total methanogenic community) after one month of fed-batch feeding of H_2_/CO_2_ but decreased to 32% after 5 months during the first stage of the enrichment. In the second stage, *Methanobacterium* dominated the methanogenic community in the inoculum (5M) but after H_2_/CO_2_ fed-batch feeding *Methanoculleus* increased in relative abundance and eventually became the dominant methanogen (T6). Previous studies reported *Methanoculleus* [25] and *Methanobacteriales* as the dominant methanogenic taxa [17] regardless of the reactor configuration. Members of the order *Methanomassiliicoccales* (class *Thermoplasmata* **Figure 2**) were present until the end of the experiment with low relative abundance, similar to a previous *ex situ* biomethanation study [26].

In reactors operated under thermophilic conditions (≥ 55°C), *Methanothermobacter* and *Methanoculleus* dominated the methanogenic community as indicated in previous studies [10,24,25,52]. Consequently, we suggest that regardless of the operational temperature, methanogens affiliated to the *Methanomicrobiaceae* and *Methanobacteriaceae* seem to be key players of *ex situ* biomethanation processes since both families dominated the methanogenic community of our enrichment culture.

The bacterial community was an integral part of the microbial community because VFA were produced and consumed (**Table 1**). Mainly acetate formation and consumption was observed, which indicates homoacetogenesis and SAO. At the end of the enrichment (T6) the dominant phylum was *Firmicutes* (91%) (**Figure S4**), which is consistent with previous findings [21], followed by the phylum *Bacteroidetes* (9%). Cooperation and competition can be expected in H_2_ biomethanation systems since the microbial community can be composed of hydrogenotrophic and acetotrophic methanogens, homoacetogenic bacteria, and syntrophic acetate-oxidizing bacteria (SAOB) [10,25] as well as chain-elongating, predatory and scavenger microorganisms. In the present study, acetotrophic methanogens were not present due to a mainly hydrogenotrophic seed sludge. In general, it is conceivable that strict acetotrophic methanogens (*Methanothrix*) coexist with homoacetogens in hydrogenotrophic communities. While a lower abundance of *Firmicutes* was described previously [10,23], a similar abundance as in this study was found by other studies [25,52,53]. In contrast, *Bacteroidetes* was found as dominant phylum by another study [54].

Throughout the enrichment, the abundance of the class *Clostridia* increased to 86% in the sixth transfer (**Figure S5**), whereas the classes *Bacteroidia* and OPB54 (*Firmicutes*) that were dominant in the inoculum decreased in relative abundance to 9% and 5%, respectively (**Figure S5**). The production of acetate from H_2_/CO_2_ (Table 1) indicated homoacetogenesis, which can be attributed to members of the *Clostridiales* that dominated the bacterial community throughout the enrichment (**Figure S6**). Some members of this order are crucial for homoacetogenesis [55] or SAO [56]. Homoacetogens and SAOB use the Wood-Ljungdahl pathway in the reductive or oxidative direction to produce acetate from CO_2_ or to oxidize acetate to CO_2_, respectively [55,57,58].

Mesophilic acetogens belong predominantly to the orders *Clostridiales* and *Selenomonadales* whereas thermophilic acetogens belong to the order *Thermoanaerobacterales* [59]. *Clostridiales and Thermoanaerobacterales* were both present in the enrichment culture (**Figure S6**) and together with the detection of acetate (**Table 1**) indicate the presence of acetogenic bacteria.

It is notable that in T6 *Lutispora* was the predominant genus with a relative abundance of 31% (**Figure S7**). The only described species of this genus is a thermophile not known to be a homoacetogen, which suggests novel homoacetogenic species may be present in the enrichment culture and supports the need to further explore such unknown microbiota. This genus belongs to the clostridial family *Gracilibacteraceae,* which was found in low relative abundance in a previous biomethanation study [26]. Other genera such as MBA03 (*Hydrogenisporales*) (19%), unclassified Family XI (*Clostridiales*) (15%), *Natronincola* (10%), unclassified *Rikenellaceae* (11%), *Fastidiosipila* (8%), *Garciella* (7%), and *Petrimonas* (5%) were also abundant parts of the bacterial community (**Figure S7**).

The acetate consumption could have occurred via SAO (e.g., by members of the classes *Synergistia* or OPB54 (*Firmicutes*) [60]) or was assimilated by acetogens and hydrogenotrophic methanogens to build biomass. The order *Thermoanaerobacterales* (class *Clostridia*) was present in lower abundance (3% in T6) with a dynamic behavior in the second stage reaching similar levels as in the inoculum. Within this order, bacteria affiliated to the genus *Gelria* (family *Thermoanaerobacteraceae*) were found. This family comprises two species known as SAOB (*Thermacetogenium phaeum* and *Tepidanaerobacter acetatoxydans*), and *Gelria* was also suggested to be involved in SAO [61]. Although the relative abundance of this genus decreased drastically towards T6, a syntrophic association with the two most dominant methanogens (*Methanoculleus* and *Methanobacterium*) of the enrichment culture is conceivable. Hence, the low concentration of acetate, especially in T6 (~123 mg L^−1^) could be explained by the activity of SAOB or high acetate assimilation by the microbial community. The SAO function of the enrichment culture might be a shared task carried out by different phylotypes since the biggest difference in abundances of *Gelria* and *Tepidanaerobacter* genera was observed at the end of T1. A previous study found *Tepidanaerobacter syntrophicus* in an *ex situ* biomethanation setup and suggested this species to be responsible for SAO [21]. Further investigations are needed by combining different omics approaches with improved isolation attempts to explore the largely unknown function of microorganisms represented only by sequence data.

### 3.3 Microbial resource management for selective production of methane

#### 3.3.1 Effect of medium composition

Our *ex situ* biomethanation experiment concomitantly enriched homoacetogenic bacteria and hydrogenotrophic methanogens similar to previous studies [10,19,24,30]. It was argued that operational conditions are crucial to shape the microbial community composition that ultimately would lead to a maximized methane yield [26]. Such control of parameters falls into the concept of microbial resource management for selective production of any desired target molecule [12,62]. Here, we showed that selective methane production with enrichment cultures that contain both homoacetogenic bacteria and hydrogenotrophic methanogens is possible by controlling the medium composition. We explored the effect of several medium components on the products of the hydrogenotrophic enrichment culture in a separate set of experiments with a focus on the products (CH_4_ and VFA) and not the microbial community. The inocula for these experiments were derived from the last culture transfer of the enrichment phase (T11).

First, we assessed the effect of yeast extract in a medium containing cysteine-HCl as reducing agent by comparing cultivation in mineral medium with (medium A) or without yeast extract (medium B). Acetate was produced up to ~5 mM in both media whereas formate was not produced at all (**Figure 4A**). However, medium A yielded 25% more biomass than medium B after the first batch-cycle feeding (p = 0.0031), even though the yeast extract concentration was as low as 0.25 mg L^−1^ (**Figure 4B**). A cultivation broth with such low concentration of yeast extract is considered as mineral medium [63].

**Figure 3.**
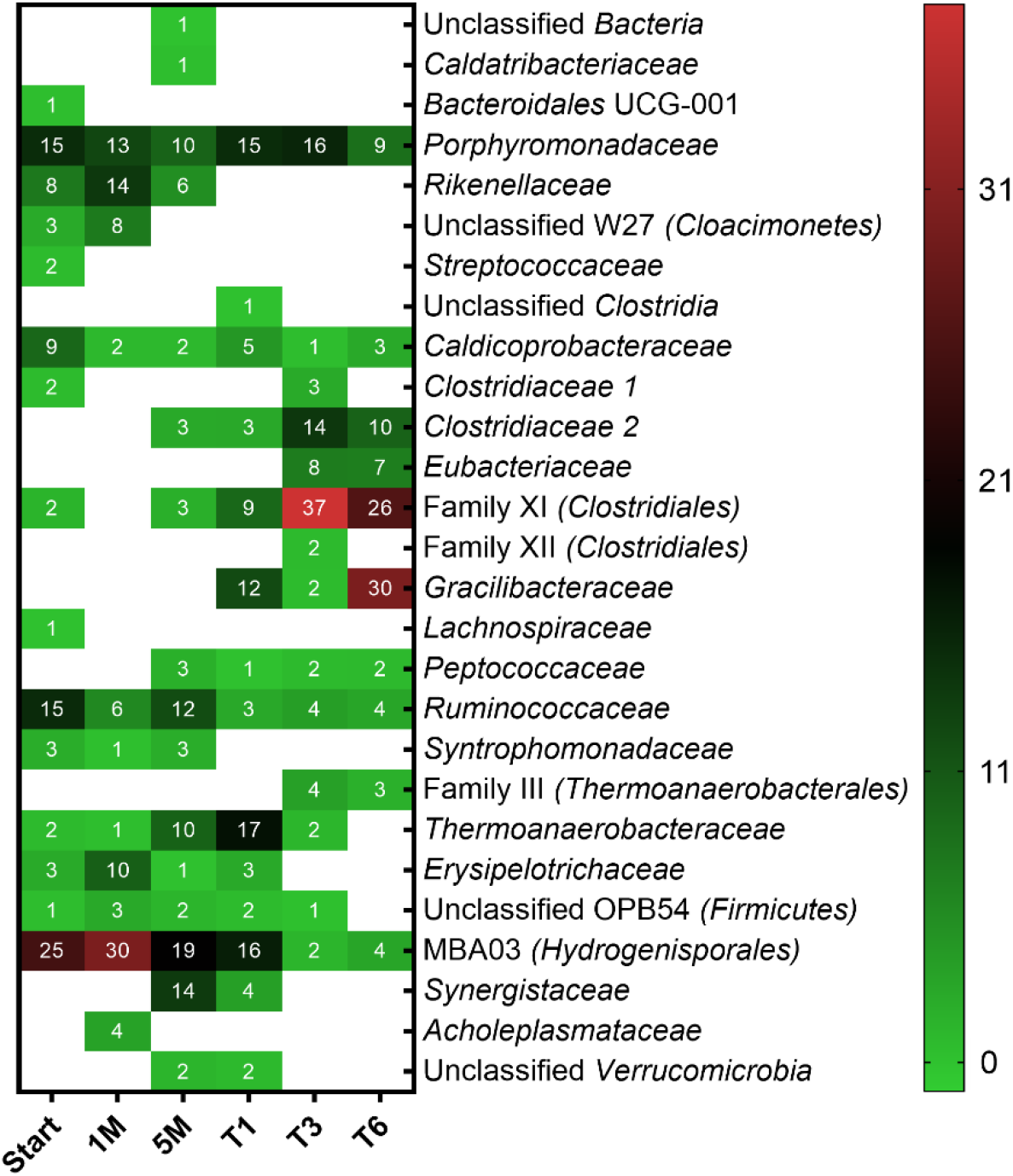
Bacterial community structure in different stages of the enrichment. Taxa with relative abundances less than 1% were filtered out from all samples. Numbers represent the relative abundance in percent and blank spaces indicate the absence of the respective taxa. Mean values of three biological replicates are presented for T1, T3 and T6 whereas single values are shown for Start, 1M and 5M.

**Figure 4.**
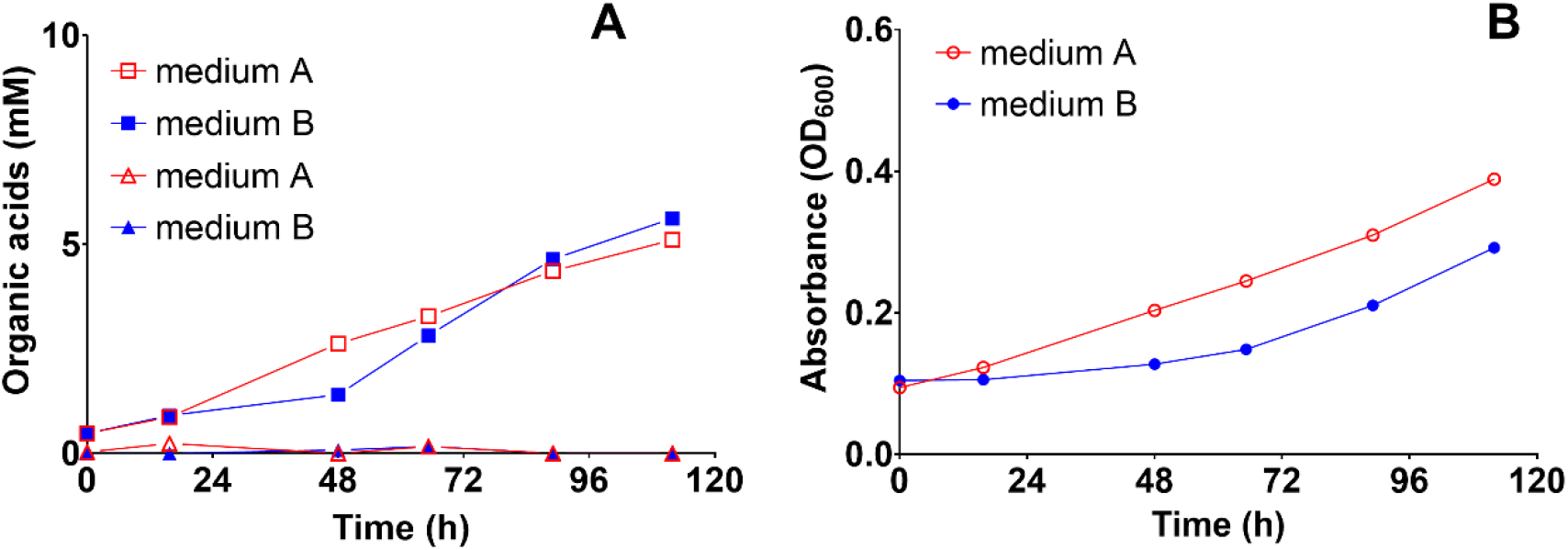
The effect of yeast extract on the production of acetic and formic acid and biomass during autotrophic feeding with H_2_/CO_2_ (80:20) in a 1-L bioreactors with medium A (containing 0.2 g L^−1^ yeast extract) and medium B (free of yeast extract but containing vitamins). Both media were reduced with cysteine-HCl. (A) Acetic acid and formic acid concentration profiles during the first batch cycle, and (B) microbial biomass growth as measured by optical density at 600 nm during the first batch cycle. The experiments were conducted in two biological replicates with orbital shaking at 200 rpm. Each data point depicts the median and range (invisible error bars are smaller than the symbol). Square: acetic acid, triangle: formic acid, circle: biomass.

In a second experiment, we tested the effect of the reducing agent on VFA formation in medium B (containing cysteine-HCl as reducing agent) and medium C (containing sodium sulfide as reducing agent). Acetate accumulated in both cultures with medium B and C but after ~300 h the concentration started to decrease and concomitantly formate concentration started to increase (**Figure 5**). This observation could indicate that formate formation resulted from acetate degradation and could be related to SAO (involving interspecies formate transfer). However, the possibility of direct formate production from H_2_/CO_2_ cannot be ruled out. The acetate concentration in medium B was as high as 20 mM (**Figure 5A)** whereas incubation for ~500 h in medium C yielded no acetate (**Figure 5B**). If cysteine-HCl is taken into account as an additional carbon source, up to 3.42 mM acetate is expected, which is far less than the accumulated acetate concentration in medium B (**Figure 5A)**, suggesting that acetate was mainly produced from H_2_/CO_2_. The final formate concentrations after 500 h were 2.4 and 2.9 mM in the cultures with medium B and C, respectively (**Figure 5**). After 500 h of operation, when the batch-cycle feeding was stopped, formate was rapidly consumed (data not shown). To our knowledge, experimental evidence of formate production in *ex situ* biomethanation has not been reported hitherto. Indeed, formate is an alternative electron donor for hydrogenotrophic methanogens [64]. Moreover, a previous study showed formate synthesis from H_2_/CO_2_ in bacteria [65]. Furthermore, pure cultures of hydrogenotrophic methanogens (*Methanobacterium formicicum)* or acetogenic bacteria transiently produced formate during H_2_/CO_2_ metabolism [66,67]. Hence, it can be inferred that homoacetogens, SAOB and hydrogenotrophic methanogens could have contributed to the concomitant formate formation along with methanogenesis. The observed formate concentration could be the result of dynamic production and consumption. The measurement of formate in micromolar concentrations is rather difficult [64]. This might explain why formate has not been reported in the liquid products in previous studies on *ex situ* biomethanation. Altogether, the results confirmed that formate was an intermediate during *ex situ* biomethanation; however, the exact mechanisms are still unclear. Reducing the sampling time intervals was important to allow formate determination in the broth. Biomass growth increased until the end of the experiment with medium B, whereas a plateau was reached after ~375 h with medium C (biomass concentration in medium B was 24% higher than in medium C). This may be explained by sulfur depletion after prolonged incubation in medium C because the medium was not replenished during the experimental period (the sulfur concentrations in medium B and C were 0.374 and 0.208 mM, respectively). The depletion of trace elements, which causes process imbalances because they are essential in enzyme complexes [68], may also explain the decreased methane production in both media after prolonged incubation. Although the biomass was less in medium C, the CH_4_ concentration was higher than in medium B (32.5%).

**Figure 5.**
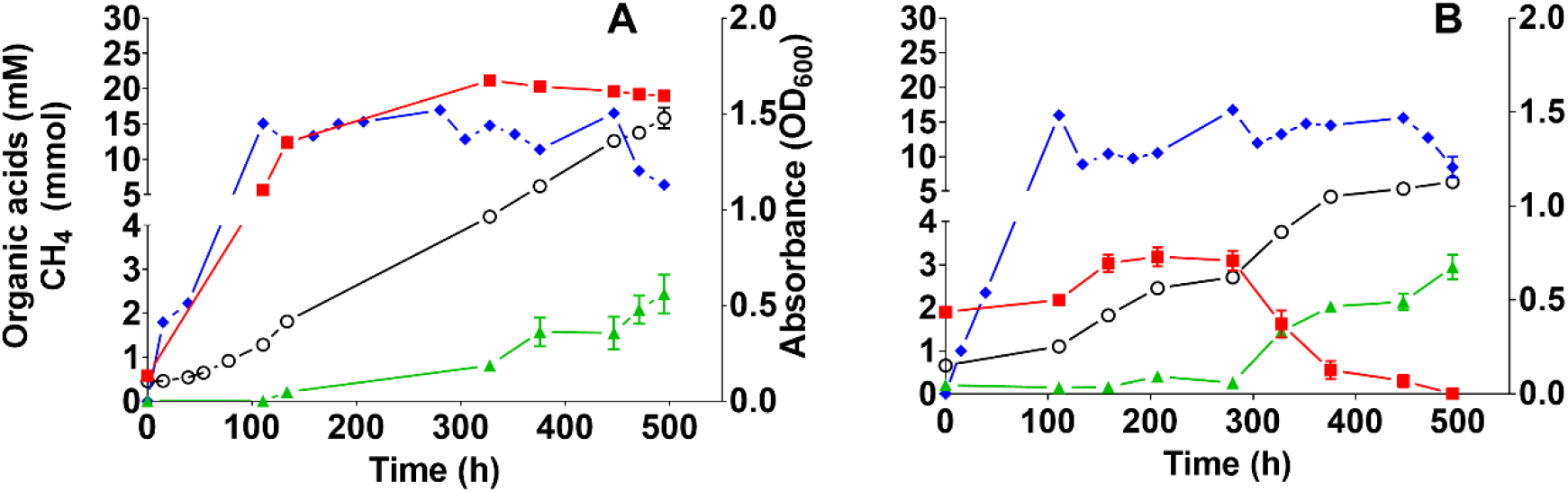
Effect of the reducing agent on the anaerobic conversion of H_2_/CO_2_ (80:20; fed-batch) in 1-L bioreactors shaking at 200 rpm. (A) Medium B (mineral medium free of yeast extract, supplemented with vitamins and reduced with cysteine-HCl), and (B) medium C (prepared as medium B but containing sodium sulfide instead of cysteine-HCl as reducing agent; see section 2.2). The experiments were conducted in two biological replicates and each data point depicts the median and range. Red square: acetic acid (mM), green triangle: formic acid (mM), blue diamond: CH_4_ (mmol), and black open circle: biomass.

#### 3.3.2 Effect of stirring intensity

Next, the effect of the stirring intensity on the methane formation rate was analyzed. Improved mixing was reported to enhance the gas mass transfer and hence methane formation rate [3,17]. As shown in **Figure 6**, the methane formation rate increased proportionally with the stirring intensity up to a maximum of ~9 mmol L^−1^ h^−1^ at 750 rpm. However, in this particle-free enrichment culture growing in a mineral medium, further increasing the stirring intensity to ≥1000 rpm was detrimental (**Figure 6**) for both methane formation and H_2_ consumption rates. Although shaking exerts a different type of mixing than stirring does, our results are in line with a previous study where shaking intensities of 200-250 rpm were already detrimental for biomethanation performed with sludge [28]. On the contrary, an *in situ* biomethanation study working with sludge reported improved gas mass transfer with a stirring intensity as high as 1000 rpm [69]. This indicates that sludge can better resist shear forces caused by high stirring intensity than enrichment cultures, so that selecting a proper mixing intensity is dependent on the type of the liquid matrix used as biocatalyst. Under optimal conditions, the enrichment culture was capable of consuming the gaseous substrate within 24 h or less, similar to times reported elsewhere [17,70]. It is to note that high shear (stirring speed ≥ 1000 rpm) may have a negative effect on the syntrophic interactions between bacteria and methanogens.

**Figure 6.**
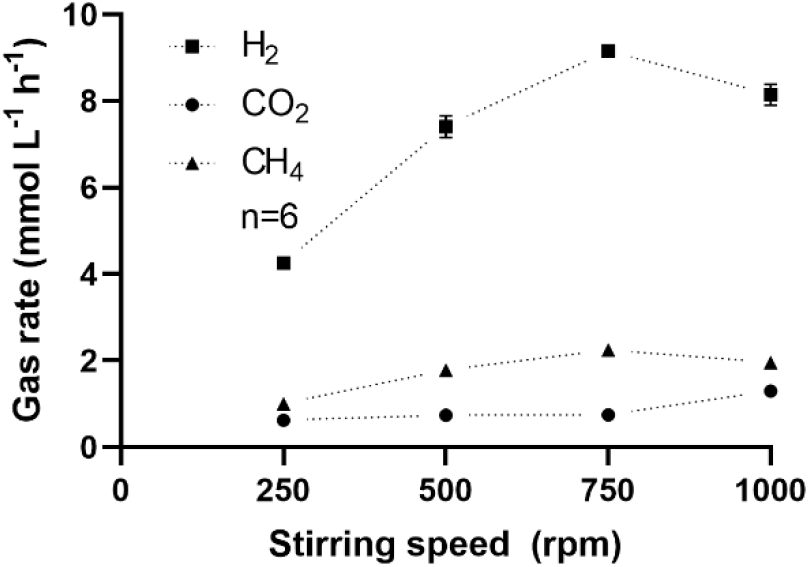
Effect of the stirring intensity on the consumption and formation rate of gases in a 1-L bioreactor with medium C. Bottles were pressurized at ~2.2 bar with a gas mixture of H_2_ (80%) and CO_2_ (20%). The rate for each stirring speed was repeated three times in duplicate biological bioreactors. Mean and standard deviation (n=6) is shown (invisible error bars are smaller than the symbol).

With an optimized medium composition (medium C) and mixing intensity (750 rpm), we obtained ≥97% methane in the gas phase, which is comparable to previous studies [1,6]. Although other measures may also affect the process performance, microbial resource management for biomethanation seems to be important when enriched mixed cultures are used as biocatalysts.

## 4. Conclusions

We showed the enrichment of a hydrogenotrophic community that successfully produced grid quality methane (≥97%) through *ex situ* biomethanation. The methanogenic community was dominated by *Methanoculleus* and *Methanobacterium*. Microbial resource management allowed the control of homoacetogenesis by directing the carbon and electron flows towards selective methane production by carefully defining the medium composition. The reducing agent played a pivotal role in controlling the production of acetate, while too high stirring intensities negatively affected *ex situ* biomethanation in a particle-free highly enriched community. Several bacterial taxa could be responsible for homoacetogenesis (mainly *Clostridia*). Thus, further investigations are needed to elucidate the physiological role of the most abundant bacterial genera in the hydrogenotrophic community.

## Supplementary Materials

Text S1: Media composition, Text S2: Chemicals and experimental operation, Text S3: Chronology of the culture transfers, Table S1. Composition of the medium component 1, Table S2. Stock solutions used to supplement the media for different experiments, Table S3. Composition of stock solutions, Table S4 Chronology of culture transfers, Figure S1 Methane concentrations during successive culture transfers in medium A, Figure S2 Correlation between optical density and biomass concentration, Figure S3 Acetate concentration profile during one batch cycle feeding with T1, Figure S4 Bacterial relative abundance at phylum level for different stages of the enrichment, Figure S5 Bacterial relative abundance at class level for different stages of the enrichment, Figure S6 Bacterial relative abundance at order level for different stages of the enrichment, Figure S7 Bacterial relative abundance at genus level for different stages of the enrichment.

## Author Contributions

Conceptualization: W.L., S.K. and M.N.; Methodology: W.L., D.P., H.S. and M.N.; Investigation: W.L.; Formal analysis: W.L. and D.P.; Data curation: W.L. and D.P.; Writing – Original Draft Preparation: W.L.; Writing – Review & Editing: D.P., S.K., H.S., H.H. and M.N.; Visualization: W.L.; Supervision: S.K., H.H. and M.N.; Project Administration: S.K. All authors have read and approved the final manuscript.

## Acknowledgments

Ute Lohse is duly acknowledged for technical assistance in library preparation for MiSeq amplicon sequencing. Cloud computing facilities used for the analysis of the amplicon were provided by the BMBF-funded de.NBI Cloud within the German Network for Bioinformatics Infrastructure (de.NBI) (031A537B, 031A533A, 031A538A, 031A533B, 031A535A, 031A537C, 031A534A, 031A532B).

