## Supplemental Material for "Microbial resource management for *ex situ* biomethanation of hydrogen at alkaline pH"

**Supplementary elements**

**Text S1.** Media composition
**Text S2.** Chemicals and experimental operation
**Table S1.** Composition of the medium component 1
**Table S2.** Stock solutions used to supplement the media for different experiments
**Table S3.** Composition of stock solutions
**Table S4.** History of the culture transfers
**Figure S1.** Methane concentrations during successive culture transfers in medium A
**Figure S2.** Correlation between optical density and biomass concentration
**Figure S3.** Acetate concentration profile during one batch-cycle feeding with T1
**Figure S4.** Bacterial community composition at the phylum level in different stages of the enrichment
**Figure S5.** Bacterial community composition at the class level in different stages of the enrichment
**Figure S6.** Bacterial community composition at the order level in different stages of the enrichment
**Figure S7.** Bacterial community composition at the genus level in different stages of the enrichment

**Text S1: Media composition**

The anoxic and sterile component 1 (**Table S1**) was supplemented with stock solutions (**Table S2**) according to the requirements of medium A, B or C. Detailed information of stock solutions is presented in **Table S3**.

**Table S1.** Composition of the medium component 1

| **Compound** | **Final concentration (g $L^{-1}$)** | | |
|---|---|---|---|
| | Medium A | Medium B | Medium C |
| $NH_4Cl$ | 0.5 | 0.5 | 0.5 |
| $KH_2PO_4$ | 0.2 | 0.2 | 0.2 |
| $MgCl_2 \times 6\ H_2O$ | 0.1 | 0.1 | 0.1 |
| KCl | 0.2 | 0.2 | 0.2 |
| NaCl | 2.0 | 2.0 | 2.0 |
| Yeast extract | 0.2 | - | - |
| Resazurin | 0.0005 | 0.0005 | 0.0005 |
| Trace elements SL10 (mL $L^{-1}$) | 1 | 1 | 1 |

Note:
Component 1 (850 mL) was made anoxic by stirring in the anaerobic chamber (97% $N_2$, 3% $H_2$) for 45 min. Sterilization was done by autoclaving at 121°C for 20 min.

**Table S2.** Stock solutions used to supplement the media for different experiments

| **Stock solutions** | **Volume added (mL $L^{-1}$)** | | |
|---|---|---|---|
| | Medium A | Medium B | Medium C |
| Selenite-tungstate solution DSM 385 (1:4 diluted) | 4 | 4 | 4 |
| $Na_2CO_3$ (29.41 g $L^{-1}$) | 34 | 34 | 34 |
| $NaHCO_3$ ( 76.00 g $L^{-1}$) | 100 | 100 | 100 |
| Cysteine-HCl (30.00 g $L^{-1}$) | 12 | 12 | - |
| $Na_2S \times 9\ H_2O$ (50.00 g $L^{-1}$) | - | - | 7.5 |
| Vitamin solution | - | 1 | 1 |

Note: Every stock solution was made anoxic by stirring in the anaerobic chamber for 30 min except for $Na_2S$ and the vitamin solution that were made anoxic with $N_2$ and sterilized by filtration.

**Table S3.** Composition of stock solutions

| **Component** | **Concentration (mg $L^{-1}$)** |
|---|---|
| Trace elements SL10 (DSMZ medium 320) | |
| $FeCl_2 \times 4\ H_2O$ | 1500 |
| $ZnCl_2$ | 70 |
| $MnCl_2 \times 4\ H_2O$ | 100 |
| $H_3BO_3$ | 6 |
| $CoCl_2 \times 6\ H_2O$ | 190 |
| $CuCl_2 \times 2\ H_2O$ | 2 |
| $NiCl_2 \times 6\ H_2O$ | 24 |
| $Na_2MoO_4 \times 2\ H_2O$ | 36 |
| Selenite-tungstate solution (DSMZ medium 385) | |
| NaOH | 500 |
| $Na_2SeO_3$ x 5 $H_2O$ | 3.0 |
| $Na_2WO_4$ x 2 $H_2O$ | 4.0 |
| Vitamin solution | |
| Biotin | 20 |
| Folic acid | 20 |
| Pyridoxine | 100 |
| Thiamine | 50 |
| Riboflavin | 50 |
| Nicotinic acid | 50 |
| Calcium pantothenate | 50 |
| Vitamin B12 | 20 |
| ***p***-Aminobenzoate | 80 |
| Lipoic acid | 50 |

Notes:

For preparing SL10, $FeCl_2$ was first dissolved in HCl (25%, 10 mL $L^{-1}$) and then diluted in water. Subsequently, other salts were added and dissolved. The 1:4 dilution was prepared with anoxic water (**Text S2**) and filter-sterilized after 30 min stirring in the anaerobic chamber.

**Text S2:** Chemicals and experimental operation

All chemicals used in this study were of the highest purity available. The gaseous substrates ($H_2$ and $CO_2$) were supplied by The Linde group (Germany). The gases were delivered via a gas mixing station equipped with mass flow controllers (Bronkhorst HIGH-TECH, Germany). The startup and each fed-batch cycle of the bottles comprised a flushing and pressurizing step. Flushing was done with a molar ratio of 4 ($H_2$) : 1 ($CO_2$) at 800 mL $min^{-1}$ ($H_2$) and 200 mL $min^{-1}$ ($CO_2$) for 10 min while pressurizing was done at half of the flow for each gas with the same molar ratio in order to reach a total pressure of ~2.2 bar inside the bottles. For flushing and pressurizing the headspace of bottles, the gas line was attached to a sterile filter (pore size 0.2 µm, Ø 25 mm; LABSOLUTE, Th. Geyer GmbH, Germany) and sterile needle (see below). To avoid pressure variation due to temperature gradients, the bottles were adjusted to room temperature prior to pressure measurement and gas sampling. Relative pressure measurements and gas sampling were done with 20 mm cannulas (Ø 0.4 mm, BRAUN) whereas flushing, pressurizing, liquid sampling and media preparation were performed with 25 mm cannulas (Ø 0.55 mm; BRAUN). The relative pressure was measured with a high-resolution manometer (LEO 5, Keller, Switzerland) attached to a sterile filter (pore size 0.2 µm, Ø 25 mm; LABSOLUTE, Th. Geyer GmbH, Germany) and cannula. Media preparation, gas and liquid sampling were done under aseptic and anoxic conditions. Prior to any media supplementation/distribution, inoculation and gas/liquid sampling the needle-syringe arrangement was flushed ten times with oxygen-free $N_2$. Prior to gas sampling, glass vials of 20 mL (LABSOLUTE, Th. Geyer GmbH, Germany) were capped with gray butyl stoppers (LABSOLUTE, Th. Geyer GmbH, Germany) and pre-flushed with argon for 20 min in a manifold arrangement with a constant gas flow controlled at ~0.2 bar.

To prepare sterile anoxic serum bottles for stock solutions and enrichment cultures, black butyl rubber stoppers and serum bottles (LABSOLUTE, Th. Geyer GmbH, Germany) were placed overnight into the anaerobic chamber (Coy Laboratory Products Inc) with an atmosphere of $N_2$ (97%), and $H_2$ (3%). 100 µL of anoxic water (see below) were added to 500-1000-mL bottles and 25 µL to 200-, 100- and 50-mL bottles. Sterilization was done by autoclaving at 120°C and 1.2 bar for 20 min.

To prepare anoxic water, 500 mL of Millipore water and a glass bar were placed in 500 mL bottles (Duran, Schott AG, Germany) and microwaved at maximum power until boiling, followed by 3 min boiling. Swirling was applied to get rid of remaining bubbles, then the water was sparged with $N_2$ in the liquid for 15 min during cooldown and the headspace was purged with $N_2$ for 15 min. $N_2$ purging was done with a constant flow and pressure controlled at 0.2 bar. The bottle was then rapidly closed with a gas-tight lid and placed in the anaerobic chamber until use.

**Table S4** History of the culture transfers for the enrichment. Since the biological replicate bottles were performing similarly, one of them was randomly selected for the next culture transfer. Letters highlighted in bold indicate the biological replicate that was used as inoculum for the next culture transfer.

| **Phase** | **Replicates** | | | |
|---|---|---|---|---|
| | ***Stage 1*** | | | |
| **Sludge** | A | **B** | C | D |
| | ***Stage 2*** | | | |
| **T1** | **A** | B | C | - |
| **T2** | A | **B** | C | - |
| **T3** | A | **B** | C | - |
| **T4** | A | B | **C** | - |
| **T5** | A | **B** | C | - |
| **T6** | A | B | **C** | - |

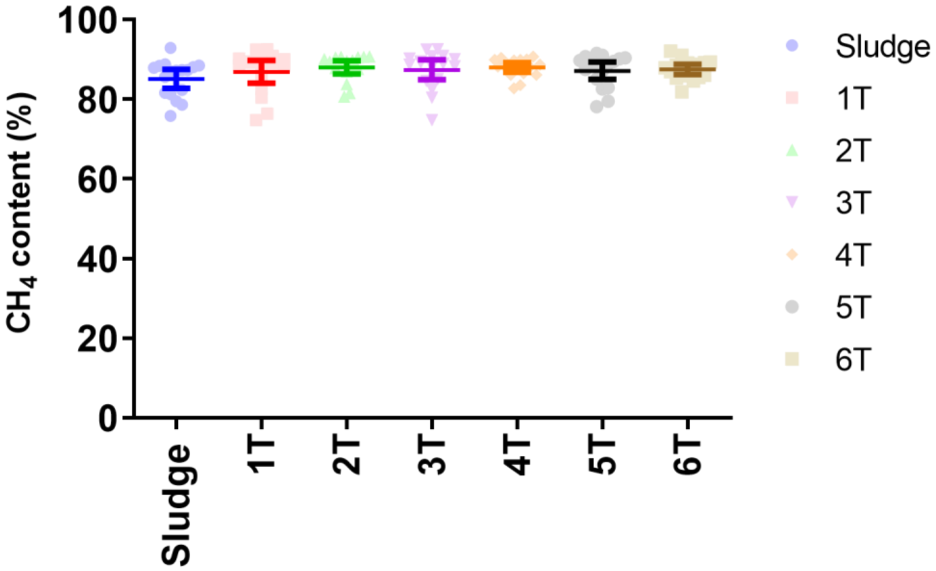


**Figure S1.** Methane concentrations during successive culture transfers in medium A. Each data point represents the methane concentration after a batch-cycle feeding of ~24 hours. The bottles were pressurized with $H_2$ (80%) and $CO_2$ (20%) to reach a total pressure of ~2.2 bar. All experiments were conducted in 200-mL serum bottles with 50 mL liquid. The inoculum size for each successive transfer was 10% (V/V) of the previous transfer. The experiments were conducted in three biological replicates for each transfer except the sludge, which had four replicates. The figure depicts the geometric mean and 95% confidence interval as well as independent measurements for each transfer.

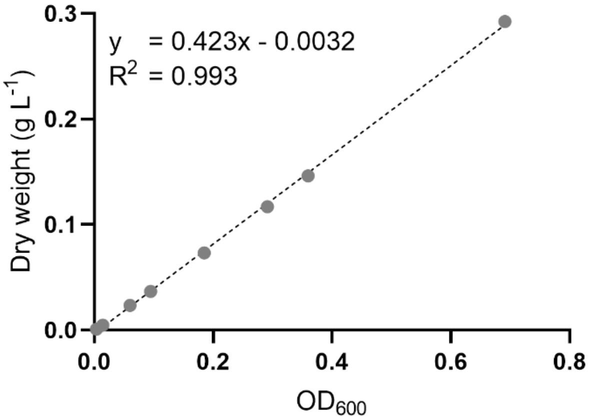


**Figure S2.** Correlation between optical density (measured at 600 nm wave length) and biomass concentration. Cultures were grown in medium A. Mean values of duplicate measurements are shown.

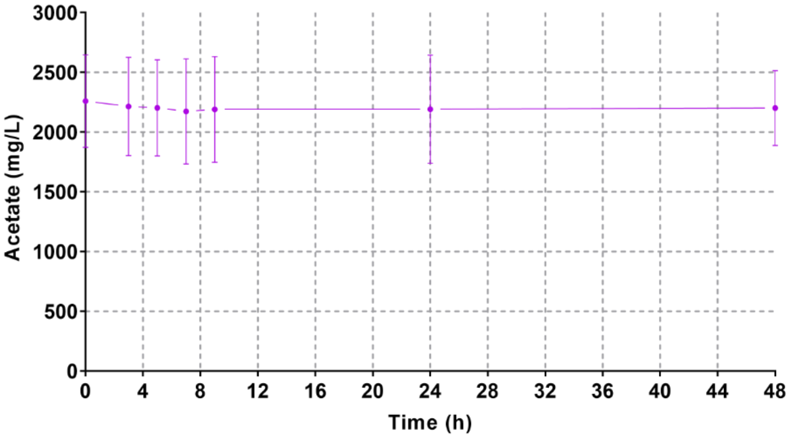


**Figure S3.** Acetate concentration profile during one batch-cycle feeding with T1. The bottles were pressurized with $H_2$ (80%) and $CO_2$ (20%) to reach a pressure of ~2.2 bar. All experiments were conducted in 200-mL serum bottles with 50 mL liquid. The figure depicts the mean and standard deviation of three biological replicates.

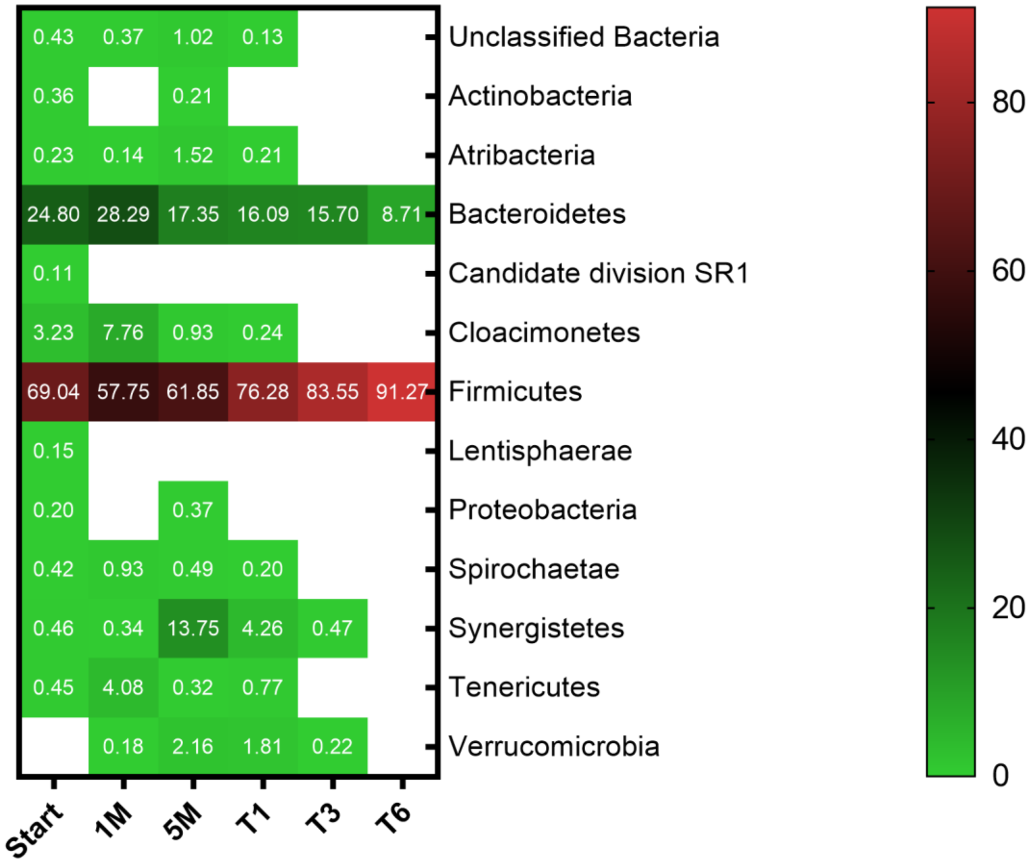


**Figure S4.** Bacterial community composition at the phylum level in different stages of the enrichment. The mean of three biological replicates is shown for T1, T3 and T6 whereas single values are shown for start, 1M and 5M. Numbers indicate the relative abundance in percentage and blank spaces indicate that the respective phylum was not detected. Phyla with a relative abundance of less than 0.01% were filtered out.

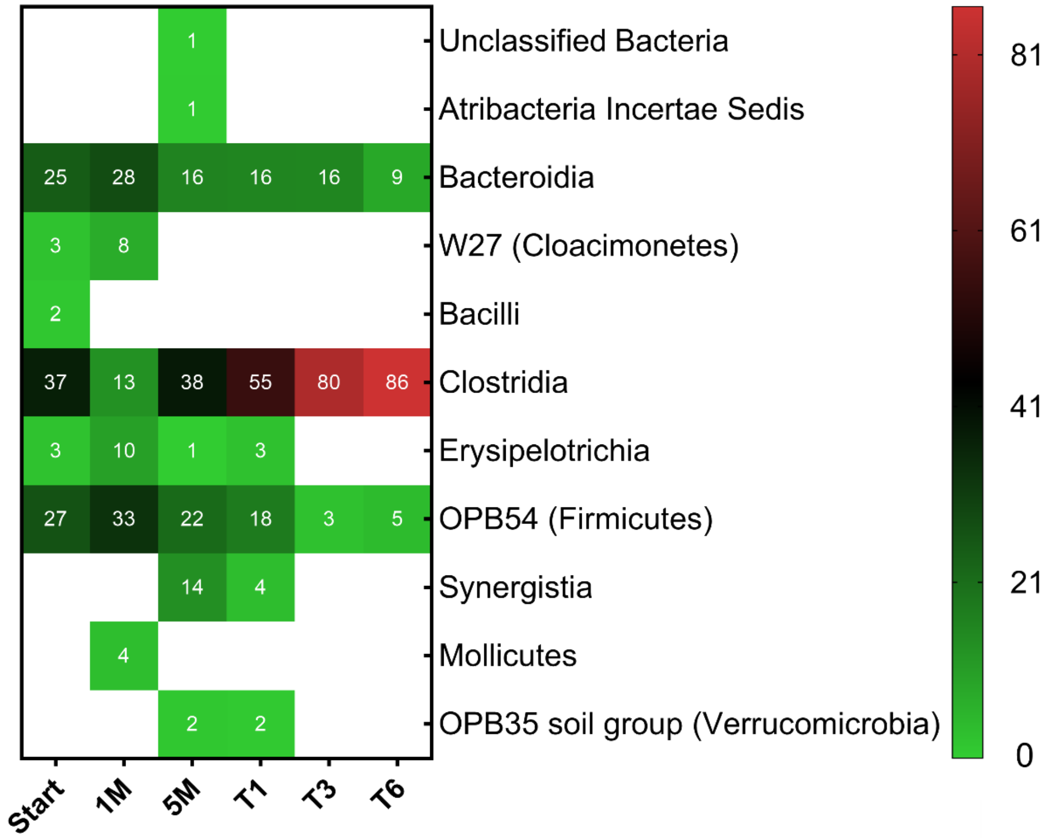


**Figure S5.** Bacterial community composition at the class level in different stages of the enrichment. The mean of three biological replicates is shown for T1, T3 and T6 whereas single values are shown for start, 1M and 5M. Numbers indicate the relative abundance in percentage and blank spaces indicate that the respective class was not detected. Classes with a relative abundance of less than 1% were filtered out.

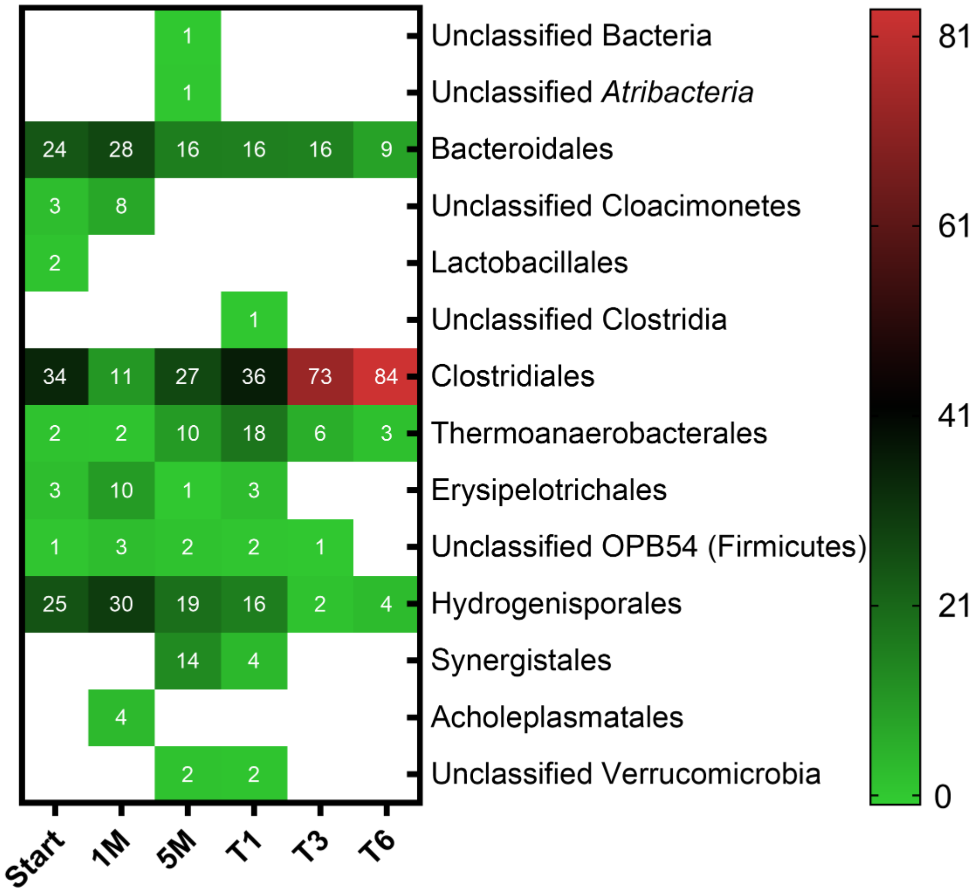


**Figure S6.** Bacterial community composition at the order level in different stages of the enrichment. The mean of three biological replicates is shown for T1, T3 and T6 whereas single values are shown for start, 1M and 5M. Numbers indicate the relative abundance in percentage and blank spaces indicate that the respective order was not detected. Orders with a relative abundance of less than 1% were filtered out.

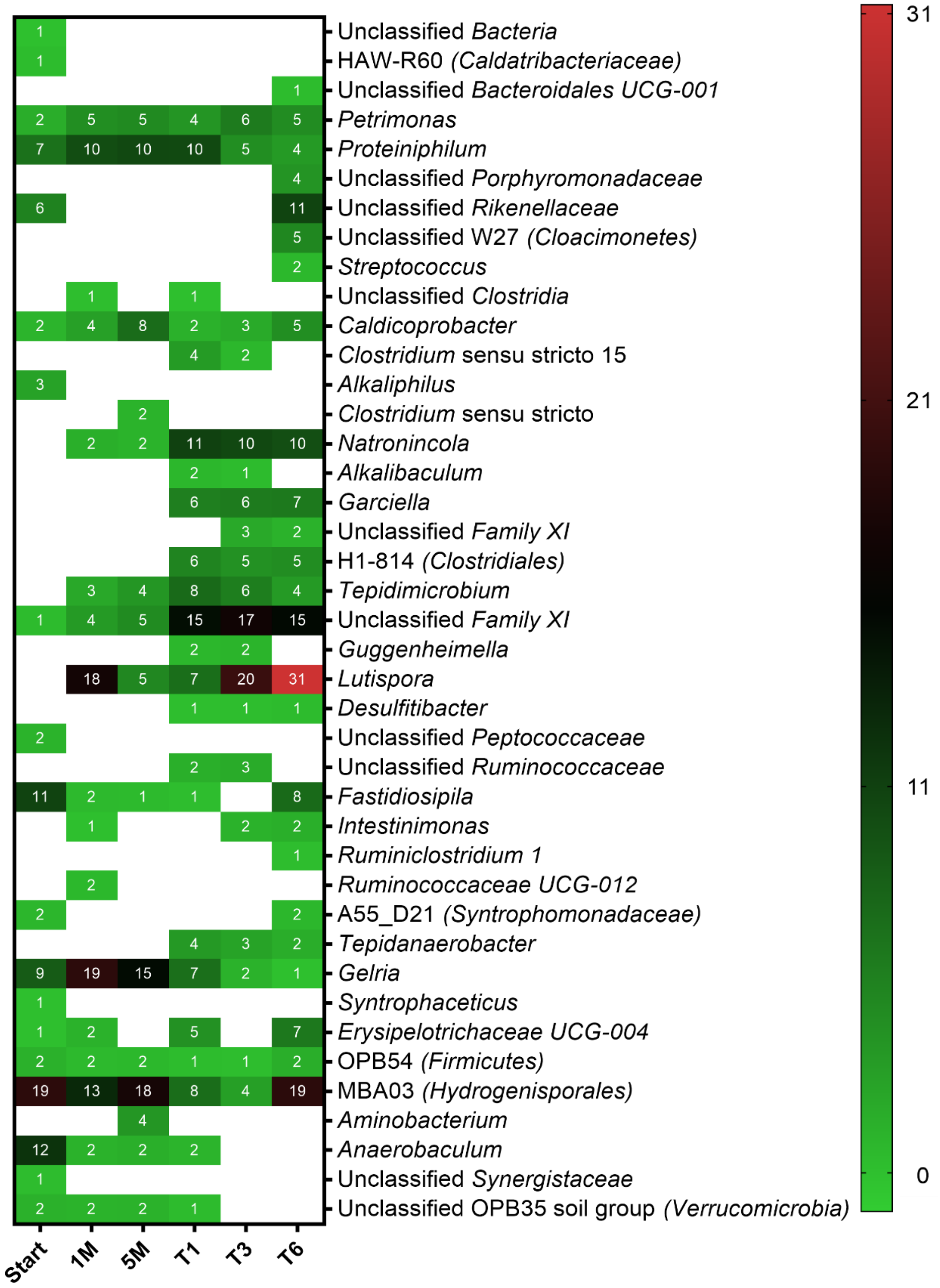


**Figure S7.** Bacterial community composition at the genus level in different stages of the enrichment. The mean of three biological replicates is shown for T1, T3 and T6 whereas single values are shown for start, 1M and 5M. Numbers indicate the relative abundance in percentage and blank spaces indicate that the respective genus was not detected. Genera with a relative abundance of less than 1% were filtered out.
